# Characterization of developmental changes in spontaneous electrical activity of superior olivary neurons before hearing onset under injectable or volatile anesthesia

**DOI:** 10.1101/2021.01.16.426864

**Authors:** Mariano Nicolás Di Guilmi, Adrián Rodríguez-Contreras

**Affiliations:** Instituto de Investigaciones en Ingeniería Genética y Biología Molecular, Dr. Héctor N. Torres, INGEBI-CONICET, Buenos Aires, Argentina, (C1428ADN); City University of New York, City College, Department of Biology, Center for Discovery and Innovation, Institute for Ultrafast Spectroscopy and Lasers. New York, New York, United States of America

**Keywords:** ketamine, Isoflurane, MNTB, in-vivo electrophysiology, multi-unit activity

## Abstract

In this work we studied the impact of two widely used anesthetics on the electrical activity of auditory brainstem neurons during development. The spontaneous electrical activity in neonate rats of either sex was analyzed under the injectable mix of ketamine/xylazine (K/X mix) and the volatile anesthetic isoflurane (ISO). We used a ventral craniotomy in mechanically ventilated pups to carry out electrophysiology recordings in the superior olivary complex (SOC) between birth (postnatal day 0, P0) and P12. To characterize neuronal activity of single and ensembles of neurons, we performed patch clamp and multi-electrode experiments under different anesthetic conditions. Our results provide the first study that compares K/X mix and ISO in the same rodent species. We demonstrate that electrical activity of SOC neurons ramps up during development, and that the firing pattern of single units recorded in K/X mix was similar to that reported in ISO anesthetized rat pups. However, ISO displayed a large scatter on its suppressing effects on electrical activity when delivered at 1.5% in the presence or the absence of K/X mix. Taken together, our results shed light on the use of anesthetics for future studies to enable electrophysiology or optical imaging studies *in-vivo* to obtain functional information on the activity of medial olivochoclear neurons and their role in auditory development.

## INTRODUCTION

The accurate organization of neuronal circuits is established during development through activity-dependent and activity-independent processes that involve the reorganization and fine-tuning of immature synaptic and cellular networks (Goodman and Shatz, 1993; Hanson and Landmesser, 2004; Kirkby et al., 2013). Before the onset of sensation, spontaneously active cells and signaling mechanisms in sensory organs have been identified as drivers of bursts of neuronal activity that are implicated in activity-dependent refinement of auditory and visual systems in different vertebrate species (Maffei and Galli-Resta, 1990; Kotak and Sanes, 1995; Lippe, 1995; Jones et al., 2007; Tritsch et al., 2007; Sonntag et al., 2009; Seabrook et al., 2017; Gribizis et al., 2019). In the auditory system of altricial rodents, calcium action potentials (APs) in cochlear inner hair cells (IHCs) initiate minibursts of APs in auditory neurons before the onset of hearing (Tritsch et al., 2007, 2010; Wang et al., 2015). During the prehearing period, IHCs are transiently innervated by direct axo-somatic efferent synaptic contacts from medial olivocochlear (MOC) neurons located in the brainstem superior olivary complex (SOC; Warr and Guinan, 1979; Simmons et al., 1996). MOC innervation is cholinergic (Glowatzki and Fuchs, 2000; Katz et al., 2004; Gómez-Casati et al., 2005), mediated by postsynaptic acetylcholine nicotinic receptors (nAChR) containing α9 andα10 subunits (Elgoyhen et al., 1994, 2001; Weisstaub et al., 2002; Lipovsek et al., 2012), and coupled to the activation of small-conductance calcium-activated SK2 potassium channels expressed in the IHCs, which mediate the hyperpolarization of IHC membrane potential in response to MOC efferent activation (Glowatzki and Fuchs, 2000). Thus, it has been proposed that MOC efferent-mediated inhibition might contribute to pattern trains of IHC calcium APs during the critical developmental period preceding hearing onset (Kros et al., 1998; Glowatzki and Fuchs, 2000; Marcotti et al., 2003; Johnson et al., 2011; Sendin et al., 2014; Moglie et al., 2018).

In recent years, different lines of genetically manipulated mouse models were used to study how modulation of the cochlear pacemaker affects the maturation of central auditory neurons and synapses. In the brainstem medial nucleus of the trapezoid body (MNTB), electrophysiological techniques were employed in slices and *in-vivo* recordings of anesthetized α9 nAChR mutant mouse pups (Clause et al., 2014; Di Guilmi et al., 2019). Despite this progress, we identified a major discrepancy in published studies based on the use of different types of anesthetics in different rodent species. The two most widely used anesthetics in brainstem studies are the injectable mix of ketamine/xylazine (K/X mix) in mice (Sonntag et al., 2009; Clause et al., 2014; Di Guilmi et al., 2019), and the volatile anesthetic isoflurane (ISO) in rats (Tritsch et al., 2010; Crins et al., 2011; Sierksma and Borst, 2017). Although different factors are taken into account to choose the type of anesthetic, few studies have addressed its potential effects on *in-vivo* spontaneous neuronal activity. For example, although previous studies have identified burst firing units in mice and rats (Sonntag et al., 2009; Tritsch et al., 2010; Clause et al., 2014), only one study in ISO-anesthetized rats reported regular firing units (Tritsch et al., 2010). This raises the possibility that the firing pattern of MNTB and other auditory brainstem neurons could be affected by the type of anesthetic used. Recent works in awake animals do not settle this issue, because optical calcium reporter fluorescence techniques were employed in auditory midbrain and cortex (Babola et al., 2018, 2020; Wang et al., 2020a), and when electrophysiology experiments were used, they were performed in the somatosensory cortex of awake mice (Wang et al., 2020b).

In this work, we studied neuronal activity with electrophysiology techniques for single unit recording and ensemble multi-unit recording in the SOC of neonate rats. Within this species, we compared the effect of K/X mix and ISO on spiking activity. We found that burst and regular firing units are present in K/X mix anesthetized rat pups, that ISO has inhibitory effects on the two types of units, and that some regular firing cells seem to be resistant to the inhibitory effects of maximal doses of ISO. Our results also demonstrate that despite the suppressive effects of anesthetics, the ensemble electrical activity of superior olivary neurons ramps up during development.

## MATERIALS AND METHODS

### Animal housing and breeding

The Institutional Animal Care and Use Committee of the City College of New York specifically reviewed and approved this study. The cohorts of adult Wistar rats used in this study were obtained from a commercial supplier at postnatal age 65 (P65, Charles River). Breeding trios of one male and two females were set for 5 days. At the completion of the breeding period, males were removed from the study while females were housed in pairs for 14 more days, and then individually until they gave birth. Pups were used from the day of birth (P0) to 12 days after birth (P12). Experiments were designed to minimize the number of animals used.

### Ventral surgery and brain processing

A total of 85 pups from 37 litters were prepared for surgery to expose the ventral skull for electrophysiology experiments. The surgical procedures used are those described by (Rodríguez-Contreras et al., 2008), with minor modifications. Neonate rat pups were initially anesthetized by an intraperitoneal injection of a mixture of ketamine hydrochloride (0.1 mg/g body weight; Ketathesia) and xylazine hydro-chloride (0.005 mg/g body weight; Anased), here referred as K/X mix. After anesthesia induction, animals were tracheotomized, intubated, and mechanically ventilated using a MiniVent type 845 ventilator (Harvard Apparatus). During surgery, anesthesia was carefully monitored on the basis of pedal reflexes, and maintained by supplementary injections of one-third of the initial dose of K/X mix as necessary (Sonntag et al., 2009). For the subset of experiments described in Fig. 2, animals were initially anesthetized inside an induction chamber with 3% isoflurane (ISO) carried in oxygen, and maintained anesthetized with 1.5% ISO carried in oxygen, delivered via a nose cone or a direct connection to the mechanical ventilator. A small ventral craniotomy (1.5 x 1.5 mm) was performed on each pup, and the brain vascular landscape constituted by the basilar artery and the anterior inferior cerebellar artery was exposed. The dura was carefully removed prior to recording. After successful recordings were obtained, animals received an overdose injection of Euthasol, and were removed from the setup to be perfused with a solution of 4% paraformaldehyde in 0.1 M phosphate buffer. Brains were removed from the skull and processed for histology to be sectioned in 60 µm thick slices. Brainstem slices were mounted onto glass slides and counterstained with DAPI prior to applying mounting medium and a glass coverslip. Brain sections were imaged with an LSM800 confocal microscope located at the CCNY imaging core facility.

**Figure 1.**
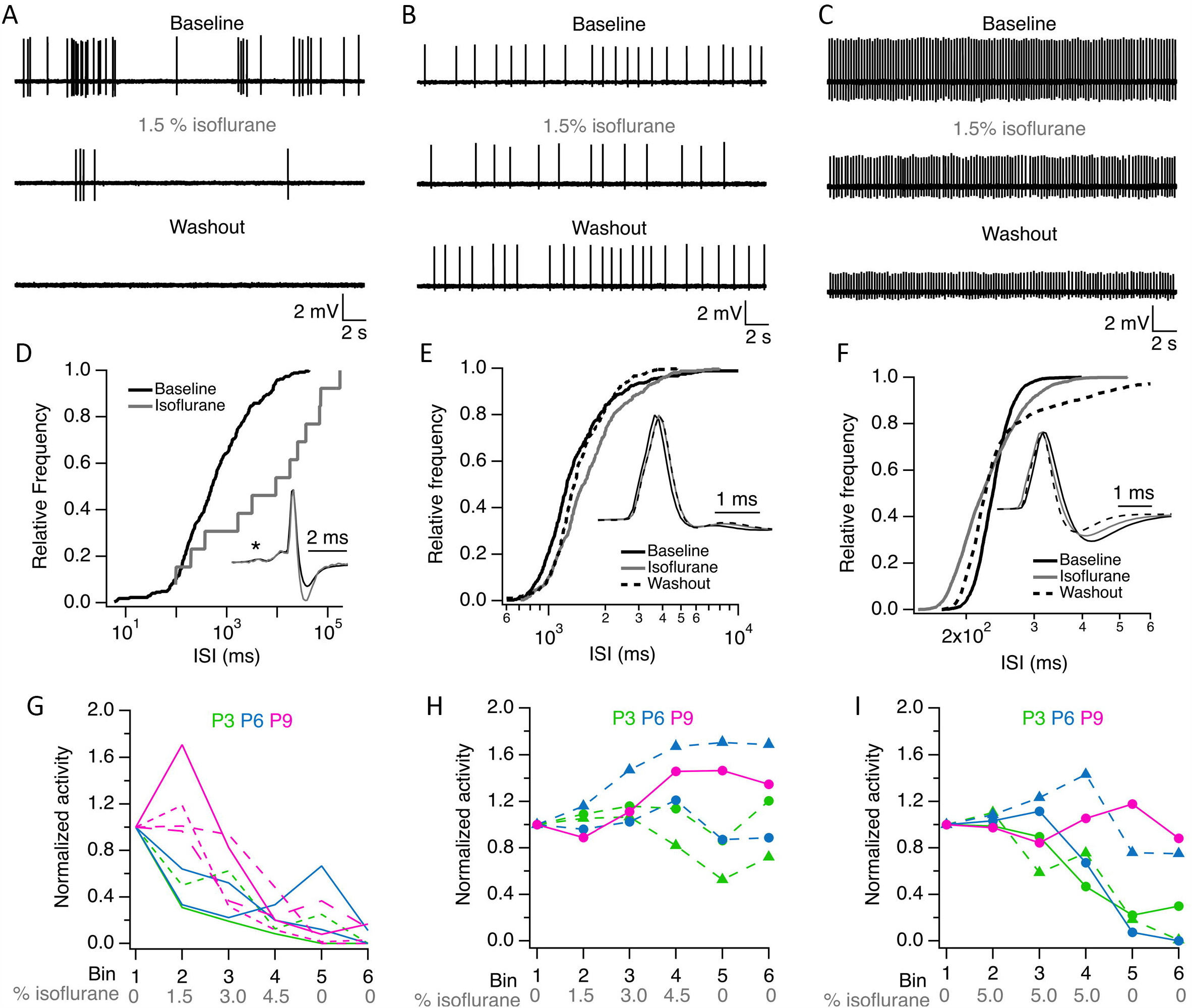
ISO inhibit single unit activity in K/X mix anesthetized pups. **A.** Burst firing unit from a P6 rat. **B**. Regular firing unit from a P3 rat. **C**. Regular firing unit from a P6 rat. **D**. Effect of ISO on the inter-spike interval (ISI) distribution of single unit shown in A. **E**. Effect of ISO on ISI distribution of single unit shown in B. **F**. Effect of isoflurane on ISI distribution of single unit shown in C. **G**. Inhibitory effect of ISO on single unit activity recorded at P3 (green), P6 (blue) and P9 (magenta). **H**. ISO resistant single units recorded at P3, P6 and P9. I. Exposure to 5% ISO inhibited 3 of 5 single units shown in panel H. ISI = inter-spike interval. Asterisk in D indicates the pre-spike in this complex waveform. **Inset** waveforms in panel D are averages of 269 and 11 action potentials in baseline and isoflurane, respectively. Inset waveforms in panel E are averages of 363, 330 and 392 action potentials in baseline, isoflurane and washout, respectively. Inset. waveforms in panel F are averages of 2538, 2599 and 2046 action potentials in baseline, isoflurane and washout, respectively. Lines and symbols in panels H and I identify the same single units under different isoflurane conditions.

**Figure 2.**
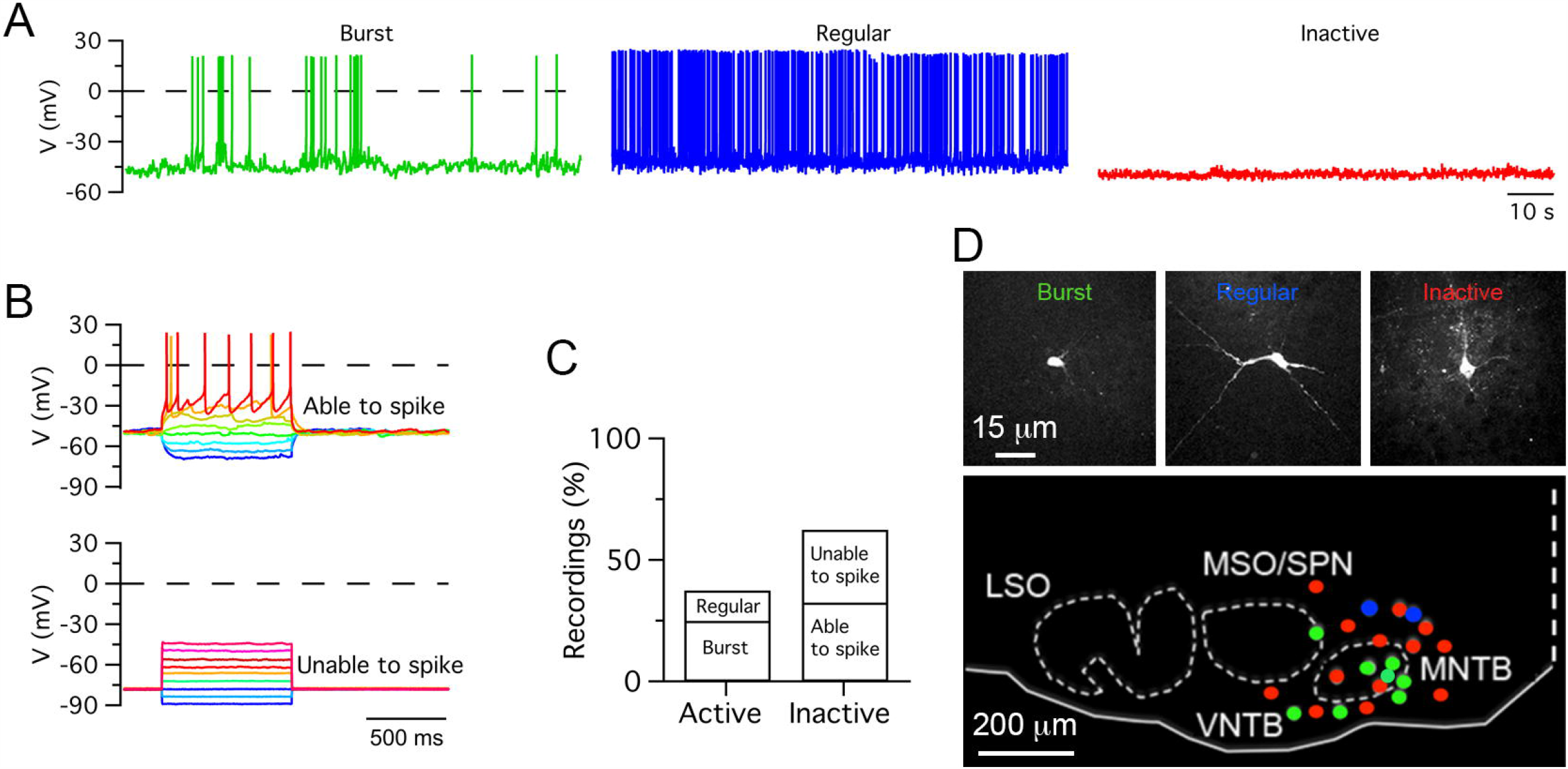
Heterogeneous electrical activity in medial olivocochlear neurons. **A.** Exemplar recordings of burst firing, regular firing and inactive cells. **B**. Inactive cells were subjected to hyperpolarizing and depolarizing voltage steps. Depolarization steps triggered action potentials in some cells but not in others. **C**. Percent of active and inactive cells. **D**. Exemplar images of cells shown in A (top row). Distribution of recorded cells in the auditory brainstem in neonate pups (bottom).

### Patch clamp recordings

Recordings were made at a depth of 200–600 µm from the pial surface using borosilicate glass pipettes (5–8 MΩ) filled with artificial cerebrospinal fluid (ACSF) containing (in mM): 125 NaCl, 5 KCl, 2 MgSO_4_, 2 CaCl_2_, 10 D-glucose, 10 HEPES, adjusted to pH 7.4 with NaOH, or intracellular solution containing (in mM): 126 K-gluconate, 20 KCl, 10 Na_2_-phosphocreatine, 4 MgATP, 0.3 Na_2_GTP, 0.5 EGTA and 10 HEPES (310 mmol.kg_-1_), adjusted to pH 7.2 with KOH. Extracellular potentials were detected in the loose-patch configuration, while intracellular potentials were recorded in the whole-cell configuration. Pipettes were advanced from the brain surface in large steps of 50 µm to a depth of 200 µm using high positive pressure (300 mbar). When in cell search mode, pressure was reduced to 30 mbar and the pipette was moved in 1.5 or 2 µm steps. The pipette tip resistance was monitored with a 10 mV test pulse in voltage clamp or when a shift in the DC was noted in current clamp. When the tip resistance increased suddenly, suggesting that a putative cell was encountered, pressure was released and slight negative pressure of 30-50 mbar was applied until a low resistance seal was formed (10-50 MΩ) for loose-patch recordings, or until a high resistance seal was formed (1-10 GΩ) for whole-cell recordings. High-resistance seals were broken by applying fast bouts of negative pressure. Voltage was recorded with pClamp9 software using an Axopatch 200B amplifier, filtered at 2–5 kHz and sampled at 20 kHz with a Digidata 1440B (MDS Analytical Technologies). Inclusion of Alexa Fluor 594 (1 %, Invitrogen) in the pipette solution allowed post hoc confirmation of recording sites or recorded cells in paraformaldehyde-fixed brainstem sections.

### Multi-electrode recordings

Prior to recording silicon probes (polytrode 4×4 16-channel arrays, NeuroNexus, A4×4–3 mm-50–125-177-A16; ∼1 MΩ) were coated with the lipophilic dye DiI (Invitrogen) for posthoc histological analysis of targeting to the medial region of the SOC. Recordings were made at a depth of 400-500 µm from the pial surface and were acquired in blocks of 10 min. All data from silicon probe recordings was sampled at 24.4 kHz, amplified and digitized in a single-head stage (TDT system III hardware, Tucker-Davis Technologies, Alachua, FL).

### Effect of isoflurane on K/X mix anesthetized pups

ISO was delivered in medical grade oxygen gas via a custom-made tube that connected a vaporizer to the animal ventilator at a rate of 1 L min^-1^. For single-unit recordings three protocols of ISO delivery were used. The first protocol consisted of acquiring 5 blocks of ten minutes each to obtain a baseline, two consecutive blocks in 1.5% isoflurane, and two washout blocks. The second was a staircase protocol that consisted of a single 10 minute-long recording where APs were quantified in six 100-second bins: baseline (0% ISO), 1.5% ISO, 3.0% ISO, 4.5 % ISO and two washout bins (0% ISO). A third protocol was used for units that showed resistance to the staircase protocol and consisted of six 100-second bins: baseline, three bins at 5% ISO, and two washout bins. The effect of ISO was evaluated in multi-electrode recordings by using a procedure similar to the first protocol described above, except that between the baseline recording and the first two blocks in 1.5% ISO, a 90 second period in 1.5% ISO was added.

### Data and statistical analyses

Unless indicated, data is presented as mean ± sem. In all cases, n indicates the number of units or cells tested, with the exception of Fig. 3 where n indicates the number of animals. Extracellular single-unit data was filtered in two steps. First, to eliminate 60 cycle electrical interference (0.6 Hz, 3 dB bandwidth); and second, with a high-pass Bessel (8-pole) filter with a −3 dB cutoff of 5 Hz using Clampfit 10.6 (Molecular Devices). All data was analyzed using NeuroMatic in Igor Pro software (WaveMetrics; Rothman and Silver, 2018). APs were detected using a positive threshold set to six times the standard deviation of the baseline noise. Single-unit recordings where an obvious gradual decrease or rundown in spike amplitude occurred were excluded from analysis. Multielectrode multi-unit data was exported as NEX files for analysis in Plexon software (Offline sorter 4.4 and Neuroexplorer). APs were detected using a negative threshold set to six times the standard deviation of the baseline noise. Multiunit activity (MUA) collected in every channel was considered. All statistical tests were performed with Statistica 7.0 software (Stat Soft; RRID:SCR_014213). Nonparametric Mann–Whitney test was used. Values of p<0.05 were considered significant.

**Figure 3.**
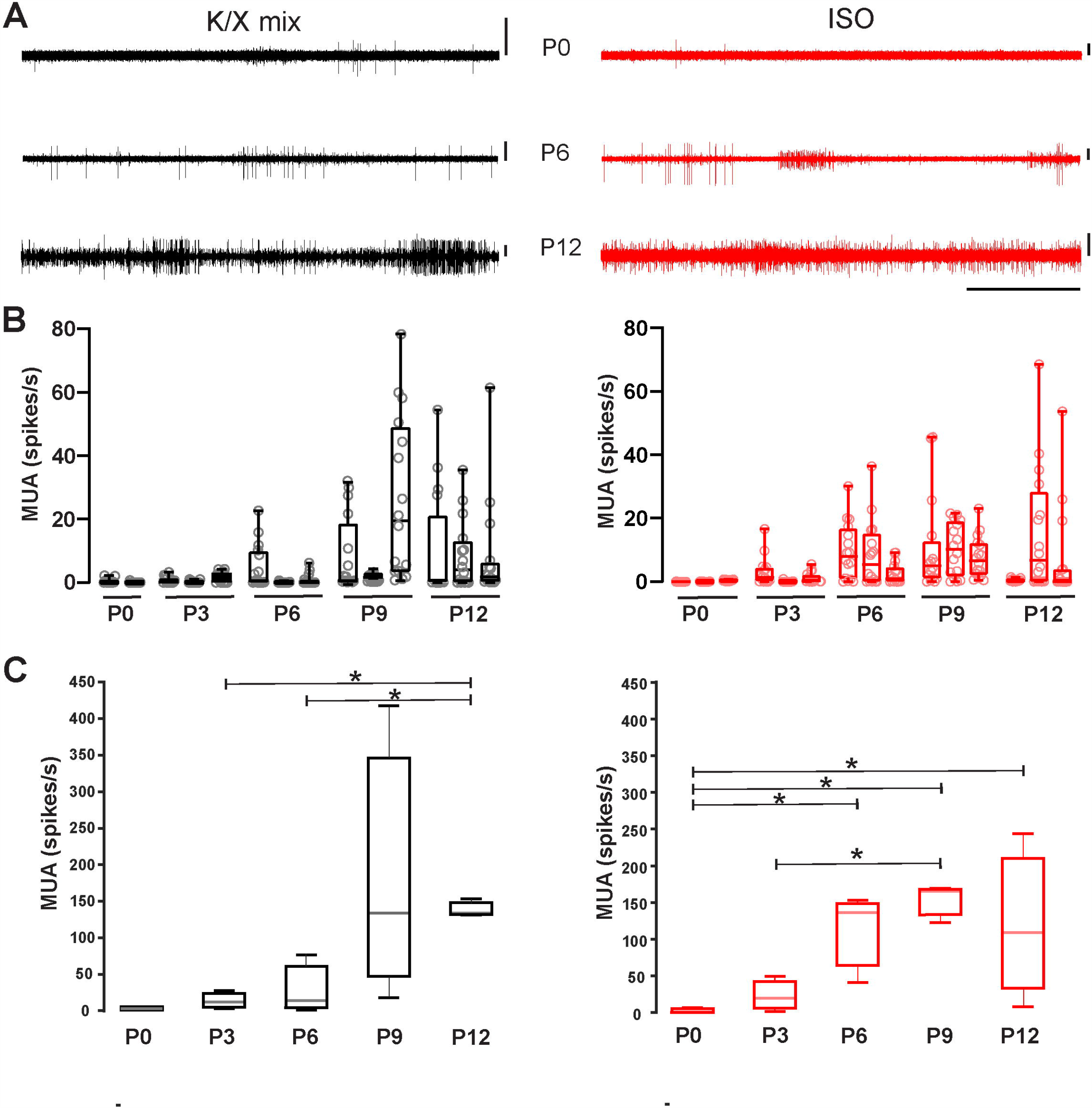
Increase in multiunit activity in the MNTB before hearing onset. **A.** Representative recordings acquired with a polytrode in the medial portion of the SOC of neonate rats. Exemplar recording at P0 (upper), P6 (middle) and P12 (bottom panel) under K/X mix (left) or ISO (right panel). Vertical scale bars = 100 µV, horizontal scale bar = 5 sec. Note the increase in the activity level along development. **B**. Box plots for different animals illustrating the MUA frequency in each electrode (empty circles). **C**. Quantification of the averaged multiunit activity per developmental stage. Boxes represent interquartile range between 25% and 75%. Whiskers indicate the minimum and maximum of all data. Inside line represents the median. Under K/X mix, activity at P3 and P6 was lower than at P12 (Mann-Whitney U Test, p= 0.048, n=3). Under ISO, P0 displays lower activity than P6 (Mann-Whitney U Test, p=0.048, n=3), P9 (Mann-Whitney U Test, p=0.048, n=3) and P12 (Mann-Whitney U Test, p=0.048, n=3). Additionally, MUA was higher at P9 than P3 (Mann-Whitney U Test, p=0.048, n=3).

### Drugs and reagents

All drugs and reagents were purchased from Sigma-Aldrich (RRID:SCR_008988).

## RESULTS

### Isoflurane inhibits burst and regular firing units in the ventral brainstem of K/X mix-anesthetized rat pups

Single-unit recordings were performed in five K/X mix-anesthetized rat pups before hearing onset, and the effects of 1.5% isoflurane were evaluated on the firing properties of six single-units (one unit from a P3 pup and five units from four P6 pups). Two of the units showed burst firing and four units showed regular firing patterns under baseline conditions (exemplar recordings are shown in Figure 1A-C). These recorded units had variable responses to ISO. The activity of the two burst firing units was inhibited to 96% and 99.6% compared to baseline and did not recover during washout. The activity of three out of the four regular firing units was partially inhibited to 34 ± 21% compared to baseline, and only in one unit recovered to baseline levels during washout. Lastly, one of the four regular firing units slightly increased its activity by 2% in ISO compared to baseline, but its activity was suppressed to 19% during washout. These inhibitory and potentiating effects of ISO on single-unit firing properties were observed in the inter-spike-interval histograms shown in Figure 1D-F. Small changes in the waveform of recorded units were also noted (insets in Figure 1D-F).

To address the variation of 1.5% ISO effects on recorded units, additional experiments were performed on thirteen single-units recorded from two P3 pups, two P6 pups, and three P9 pups by increasing the ISO dose from 0% to 4.5% in 1.5% steps. It was found that two P3 units, two P6 units and four P9 units showed clear signs of dose-dependent inhibition by ISO compared to baseline. These cells did not recover during the washout period (Figure 1G). In a total of five single-units, evidence of potentiation or resistance to increased doses of ISO was found (Figure 1H). Three of these units were inhibited by prolonged exposure to 5% ISO, while two units from P6 and P9 pups showed signs of resistance and potentiation even at such high dosage (Figure 1I).

Altogether, these experiments show that burst and regular firing cells are present in K/X mix-anesthetized rat pups, that most burst and regular firing units are inhibited permanently by ISO, while some regular firing units appear to be resistant to inhibition and show indications of potentiation after ISO is delivered to K/X mix-anesthetized rat pups. Resistance and potentiation to ISO was stronger in P6 and P9 rat pups.

### Heterogeneous population of single-units: inactive *versus* active neurons

It was noted that from a total of nineteen recorded single-units, five units fired in a burst type pattern, while the rest fired in a regular spike pattern. Furthermore, three of the burst firing units did not show complex waveforms adjudicated to MNTB neurons. These observations raised the possibility that other SOC neurons may fire burst type or regular type patterns. One caveat of loose-seal patch clamp recordings is that cells cannot be identified, unless iontophoretic methods are used to label them (Pinault, 1996; Cid and de la Prida, 2019). As an alternative to identify the morphology and location of recorded cells, whole-cell current clamp recordings were performed in forty ISO anesthetized pups between ages P0 and P11 (Figure 2). Active and inactive cells were identified in current clamp recordings. Active cells fired APs in burst (n=10) or regular type (n=5) patterns, while inactive cells did not fire APs at rest (n=25; Figure 2A). Two kinds of inactive cells were further identified based on their response to depolarizing current injection. Some cells fired APs (n=13), while other cells did not (n=12; Figure 2B). Therefore, inactive cells were classified as able or unable to fire APs (Figure 2C). After histological processing, the morphology and location of twenty-three recorded cells was confirmed in the medial region of the SOC (Figure 2D). In sum, these experiments show that ISO-anesthetized rat pups have active and inactive neurons in the SOC. Several cells located in the medial region of the SOC fire APs in burst or regular patterns. Some inactive cells are capable of firing action potentials upon depolarization of their membrane potential, while others cannot fire action potentials suggesting irreversible inhibition or a different cellular phenotype (i.e., non-neuronal glial).

### Multiunit activity increases during postnatal development

Next, the effect of the injectable and volatile anesthetics on the ensemble activity of the medial SOC was evaluated with multi-electrode probes in pups aged between P0 and P12 (n=29; Figure 3A). Despite the large variability of spontaneous MUA activity within each developmental group, the overall MNTB spontaneous activity ramped up during development under K/M mix (n=14; Figure 3B-C, left panel) or ISO (n=15; Fig. 3B-C right panel). It is important to note that the mean firing rate was similar under both conditions. Statistical analyisis showed that under K/X mix, the MUA activity was significantly lower at P3 compared to P12 (Mann-Whitney U Test, p= 0.048, n=3), and at P6 compared to P12 (Mann-Whitney U Test, p= 0.048, n=3). Under ISO, P0 displayed lower activity compared to P6 (Mann-Whitney U Test, p=0.048, n=3), P9 (Mann-Whitney U Test, p=0.048, n=3) and P12 (Mann-Whitney U Test, p=0.048, n=3). Additionally, MUA was higher at P9 than at P3 (Mann-Whitney U Test, p=0.048, n=3). Taken together, these results indicate that the overall level of spontaneous electrical activity increases during postnatal development and that the maximal ensemble firing rate was similar between the two anesthetics.

### Developmental profile of ISO inhibition

To examine whether ISO may act over the anesthetic effect of K/M mix and considering that both anesthetics have different synaptic targets (MacDonald et al., 1987; Wang et al., 2020b), we implemented the recording protocol illustrated in Figure 4A. Figure 4B displays three examples of MUA activity at different developmental ages (P3, P6 and P9) along the experimental protocol. In all cases, application of 1.5% ISO depressed the spiking activity, and a partial or total washout could be observed. The activity level was quantified after application of 1.5% ISO relative to K/X mix baseline (Figure 4C). Grouping data into two age ranges showed that ISO had a greater initial inhibition at P0-P6 in comparison with P9-P12 (Mann-Whitney U Test p=0.002, n=3). However, after 20 minutes ISO inhibited at all ages. In conjunction with loose-seal patch clamp recordings (Figure 1), these experiments show that the inhibitory effect of ISO is heterogeneous before hearing onset.

**Figure 4.**
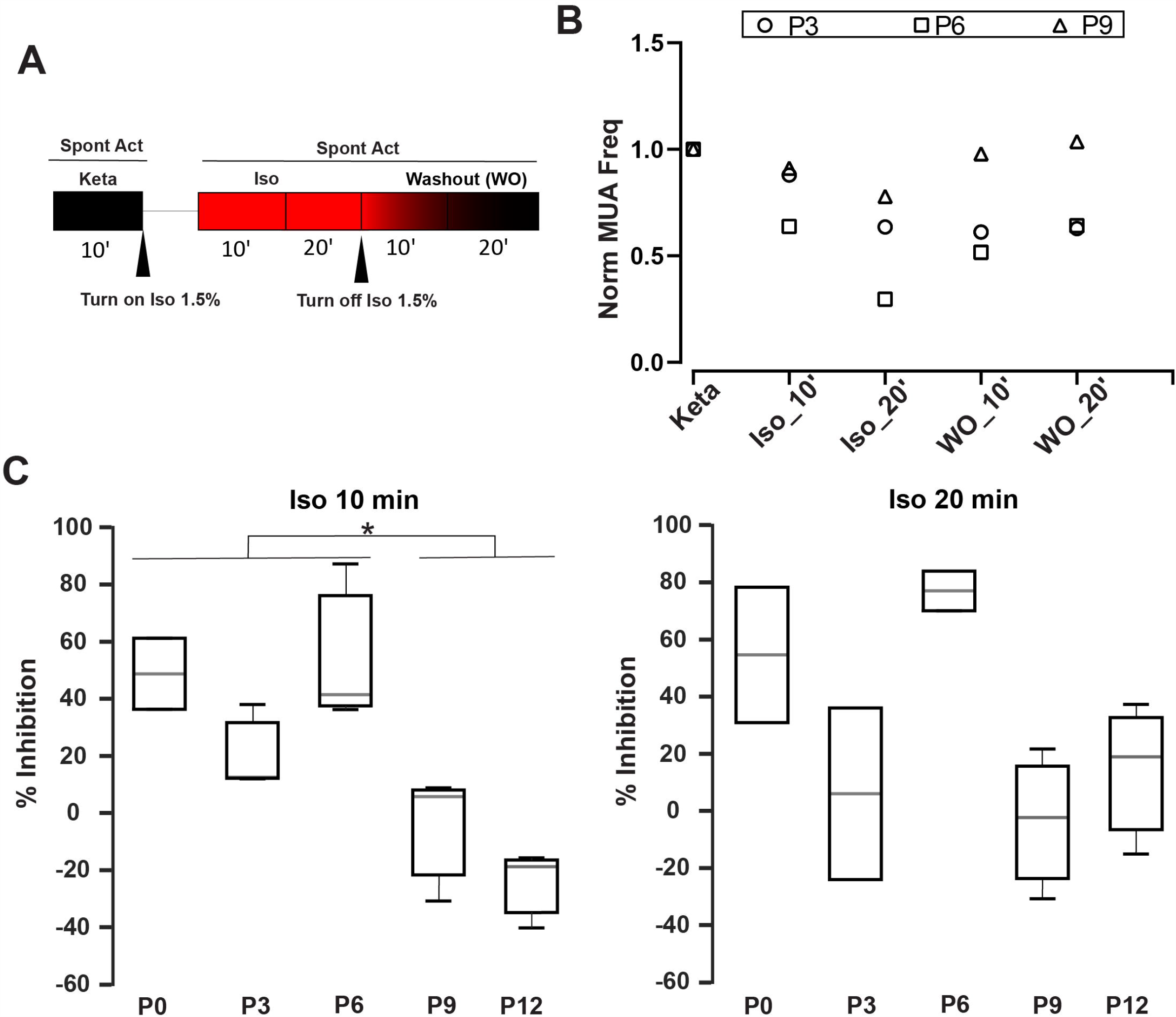
Time-dependent inhibition of Isoflurane. **A.** Schematic diagram of the pharmacological protocol for testing Isoflurane inhibition. The first step was to record the spontaneous activity during 10 min under K/X mix (black box). Then we applied Isoflurane 1.5% for 90 seconds followed by two recording rounds of 10 min to obtain MUA after 10 or 20 min respectively (red boxes). After that, ISO was turned off to evaluate the washout for the next 20 min. **B**. Normalized MUA frequency relative to the K/X mix state (Norm MUA freq) at different developmental ages (P3, P6 and P9). **C**. Percentage of Inhibition (MUA frequency under ISO / MUA frequency under K/X mix) after 10 (left) or 20 minutes (right) with ISO at different developmental stages. Grouping stages from P0 to P6 displayed a significant inhibition in comparison with stages P9-P12 (Mann-Whitney U Test p=0.002, n=3).

## DISCUSSION

In this work we decided to study the impact of K/X mix and ISO, the two most widely used anesthetics in auditory developmental studies of spontaneous electrical activity in neonate rodents. General anesthetics are widely used for surgical procedures in animal models and sometimes are required during *in-vivo* experiments. For example, recent studies have targeted dorsally accessible brain structures like the inferior colliculus (IC) and auditory cortex (AC) in awake transgenic mice that had been exposed to anesthetics during preparatory surgery (Babola et al., 2018, 2020). However, if ventrally accessible brainstem structures are targeted to perform imaging and/or electrophysiological recordings, a general anesthetic must be used to perform recordings in animals that are mechanically ventilated (Rodríguez-Contreras et al., 2014; Di Guilmi et al., 2019), or that are immobilized to facilitate accurate electrode targeting (Sonntag et al., 2009; Clause et al., 2014).

This study provides three main findings that are relevant to understand the role of MOC neurons and other brainstem neurons in auditory system development. First, to the best of our knowledge, this is the first study that compares K/X mix and ISO in the same rodent species. The firing pattern of single units recorded in K/X mix was similar to that reported in ISO anesthetized rat pups (Tritsch et al., 2010). Furthermore, this is the first study that identified burst-firing neurons outside the MNTB, which due to their location were mapped to the medial portion of the superior olivary complex. MOC neuron somata are diffusely located in the ventral nucleus of the trapezoid body (VNTB) (Warr, 1975; Warr and Guinan, 1979; Guinan et al., 1983) and are difficult to identify without labeling on slices (Cadenas et al., 2020). We consider that our results shed light on the use of anesthetics for future studies to enable patch-clamp electrophysiology combined with imaging studies on MOC neurons *in-vivo*. Second, it was found that ISO displayed a large scatter on its suppressing effects on electrical activity of auditory brainstem neurons. First, in pups that were pre-anesthetized with K/X mix, around two thirds of recorded cells were inhibited by 1.5% isoflurane, the isoflurane EC50 estimated under controlled conditions (Wang et al., 2020b). However, an estimated one third of recorded units were resistant to 1.5% and were inhibited by 5% isoflurane delivered over 20 minutes. Second, burst and regular firing units were observed in pups purely anesthetized with 1.5% ISO. Inhaled anesthetics diffuse from the alveoli into arterial and capillary blood, and are assumed to equilibrate rapidly with the well-perfused central nervous system (Hemmings et al., 2005). Thus, the heterogeneity of inhibitory effects observed in the results of this study, may be explained by the hypothesis that the physical and functional properties of the vasculature, the diffusive properties of the extracellular space between vessels and neurons, and the clearance mechanisms of extracellular fluid available during this developmental window may be important factors that determine the access of anesthetics to target neurons in the SOC (Nicholson et al., 2000; Shi and Rodríguez-Contreras, 2016; Mestre et al., 2020). One caveat of the present study is that we were not able to study the effect of different concentrations of K/X mix on auditory brainstem neuronal activity. Since the K/X mix and ISO have different synaptic targets (i.e. postsynaptic NMDA receptors (MacDonald et al., 1987) or presynaptic effect (Wang et al., 2020b), respectively), we evaluated the ISO inhibition under K/X mix. In agreement with previous reports, depressant percentage depended on ISO concentration and the developmental age in the MNTB (Wu et al., 2004; Wang et al., 2020b). Different reasons related to the plastic changes on these synapses (Wu et al., 2004) may support these results. Furthermore, we observed that a group of recorded cells were inactive. One of the responsible factors of neuronal excitability is the resting membrane potential (Vm). Volatile anesthetics induced negative potentials on cortical cells (Petersen et al., 2003) and the hyperpolarization of postsynaptic MNTB neurons in brain slices (Wang et al., 2020b). Rusu and Borst (2010) demonstrated in slices containing the MNTB that Vm decreases during the first postnatal week becoming progressively more negative. During the first two postnatal weeks, the heterogeneity in the intrinsic properties maturation is larger than in adulthood, and probably it is a substrate of our observations. However, the effect of ISO on the Vm of these cells is under debate (Wu et al., 2004; Wang et al., 2020b) and a further study of molecular mechanisms during development is necessary to clarify the results. The third important finding in this work was that electrical ensemble activity of superior olivary neurons ramps up during development before hearing onset. Note that the ensemble activity displayed the same developmental profile under both anesthetics. A recent report describes that both *in-slice* and *in-vivo*, ISO preferentially inhibits spike transmission (Wang et al., 2020b). In that work where the authors proposed ISO as a low-pass filter, the ISO mechanism of action was studied on MNTB slices but the *in-vivo* effect was examined in a cortical structure. In the present paper, we demonstrated that at the brainstem level, ISO alone does not affect significantly the firing rate of MNTB neurons. This result is promising in terms of future studies specially for those where a ventral procedure is required to perform imaging and/or electrophysiological recordings. Due to the little influence on metabolism and its simple handling, specially for long duration experiments (Albrecht et al., 2014), ISO displayed an operative advantage over K/X mix.

## Author Contributions

Designed and performed: A.R.-C (juxtacellular & multielectrodes), M.N.D.G. (multielectrodes); Analyzed data: M.N.D.G. & A.R.-C; Wrote the first draft of the paper: M.N.D.G. & A.R.-C.; Edited the paper: A.R.-C; Contributed unpublished reagents/analytic tools: A.R.-C.

## Acknowledgments

We would like to acknowledge the Science Division Core Imaging Facility and Dr. Jonathan Levitt for providing access to a microtome to conduct histology procedures.

